# Gene families with stochastic exclusive gene choice underlie cell adhesion in mammalian cells

**DOI:** 10.1101/2020.08.24.264747

**Authors:** Mikhail Iakovlev, Simone Faravelli, Attila Becskei

**Author notes:** These authors contributed equally (Mikhail Iakovlev, Simone Faravelli).

## Abstract

Exclusive stochastic gene choice combines precision with diversity. This regulation enables most T-cells to express exactly one T-cell receptor isoform chosen from a large repertoire, and to react precisely against diverse antigens. Some cells express two receptor isoforms, revealing the stochastic nature of this process. A similar regulation of odorant receptors and protocadherins enable cells to recognize odors and confer individuality to cells in neuronal interaction networks, respectively. We explored whether genes in other families are expressed exclusively by analyzing single cell RNA-seq data with a simple metric. Chromosomal segments and families are more likely to express genes concurrently than exclusively, possibly due to the evolutionary and biophysical aspects of shared regulation. Nonetheless, gene families with exclusive gene choice were detected in multiple cell types, most of them are membrane proteins involved in ion transport and cell adhesion, suggesting the coordination of these two functions. Thus, stochastic exclusive expression extends beyond the prototypical families, permitting precision in gene choice to be combined with the diversity of intercellular interactions.

## INTRODUCTION

The combinatorial principle plays an important role in the evolution of complex organisms. A large proportion of the mammalian genomes encodes regulators, especially transcription factors (1), which determine what combination of genes will be turned on and off. Each cell type expresses a distinct set of genes, a form of phenotypic diversity that has been studied by single cell expression profiling, such as single-cell RNA-seq, with an unprecedented throughput. These and other single cell studies revealed that the combinations of expressed genes are not uniform in a cell population but are subject to substantial variations, in part due to the inherently stochastic nature of RNA synthesis and degradation (2-4). Notwithstanding this variability, specific combinatorial patterns have been identified, among which the stochastic exclusive gene expression of the odorant and T-cell receptors has received a widespread attention.

Each olfactory neuron expresses a single odorant receptor isoform upon a random choice from more than thousands of gene isoforms (5, 6) and elicits a signal in response to a specific odor. Thus, precision is combined with diversity. A similar principle underlies the immune response: each lymphocyte expresses a single antigen receptor randomly chosen from a large repertoire. The receptor isoforms are diversified, in part, due to the stochastic gene choice of the variable domain. With the in-depth study of these systems, it became apparent that a non-negligible proportion of cells expresses more than one, typically two gene isoforms (7). These cells with dual T cell receptors may enhance the antiviral response but can also underlie autoimmune disorders (8, 9). Thus, the combinatorial stochastic choice has clear physiological implications.

A slightly different form of exclusivity was observed in the protocadherin (Pcdh) array, which encodes multi-subunit membrane proteins mediating cell-to-cell interactions between neurons (10). In this array, most cells express at least two distinct variable α-isoforms from a repertoire of 12 genes, one from the paternal, one from the maternal chromosome (11), which is highly reminiscent of the cells with the dual T-cell receptors. These findings indicate that the strict definition of exclusivity – one gene (isoform) per single cell – needs extending to account for the observed distributions and for averages greater than one.

These observations lead to the question about how to define exclusive expression in terms of a probability distribution. Is the expression of T-cell receptor isoforms exclusive if cells with dual T-cell receptors constitute 1%, 50% or 90% of the population? What if three different receptor isoforms were to be expressed in some of the cells (12)? Recently, the degree of exclusivity in the stochastic gene choice of the Pcdh gene array was quantified with a probabilistic approach that defines exclusivity independently of the mean number of expressed genes in an array (13). This definition of stochastic exclusivity implies that the distribution of the number of expressed gene isoforms is narrower than expected from the purely random, independent expression of the genes in the array. Thus, stochastic exclusivity reflects simply the precision in gene choice irrespective of the underlying mechanism, let it be chromosomal looping during gene activation, negative feedback or allelic exclusion after DNA recombination. As an example, precision implies that the majority of the cells express three gene isoforms and the frequency of the cells expressing fewer or more than three isoforms is low in comparison to a distribution resulting from a random independent choice with equal mean.

Here, we examined single-cell RNA-seq data to assess the presence of stochastic exclusive gene choice in chromosomal segments and gene families.

## RESULTS

### Dichotomization of RNA-seq counts

We analyzed RNA-seq datasets consisting of at least 100 single cell measurements of a well-defined cell type isolated from the mouse Mus musculus (Table S1). Neurons from various locations in the nervous system were included, such as somatosensory neurons from dorsal root ganglions (14), dopaminergic neurons (15) and corticostriatal neurons from the visual cortex (16) (Table 1). Non-neuronal cell types encompassed nearly all organs: two types of lymphocytes, CD8+ T-cells (17) and type 17 helper cells (Th17) (18); dendritic cells from the bone marrow (19), cardiomyocytes (20), endothelial cells (21), enterocytes (22), fibroblasts (23), kidney duct cells (24), thymus epithelial cells (25), prostate stromal cells (26), type I and II alveolar cells from the lung (27); hepatoblasts and hepatocytes from the liver (28), pancreatic endocrine cells (29). Undifferentiated cell types were represented by embryonic stem cells isolated from embryos (30) and ES cell cultures (31).

Combinatorial gene choice is typically quantified for genes whose expression is characterized by two states, variously named as ON/OFF, binary or bimodal expression. A large fraction of genes displays bimodal expression (Figure S1A) (32). To determine the proportion of ON and OFF cells, the single-cell RNA counts have to be dichotomized. For this purpose, we compared two classes of methods. In the moment-based method, the averages or variances of the full or part of the distributions are calculated. The second method relies on the fitting of probability mass or density functions (pdf). The moment based methods are more robust but lack a uniform mathematical framework (Figure S1B-D). Conversely, the pdfs have mathematically well-defined dichotomization points but their fitting is less robust, especially when there are few cells in the OFF or ON expression states or when the measurement error is larger. In order to combine the advantages of the two approaches, we aimed at selecting the moment based approach that correlates the most with the dichotomization using pdfs.

We tested three types of moment-based methods: the Variance Reduction Score (VRS), Fraction of Maximal values (FM) and Geometric Trimmed Mid-Extreme threshold (GTME) (Materials and Methods). The VRS quantifies the extent to which a given threshold reduces the sum of the variances of the two subpopulations relative to the unsplit population (33). The threshold minimizing the VRS was selected for the dichotomization. We devised two additional methods based on biological control principles, the FM and GTME. The FM is based on the assumption that a biological function can be performed as long as a variable in the ON state does not deviate too much from an optimal level. Accordingly, we defined the FM-threshold as the one tenth of the observed maximal values in the distribution. The GTME threshold is the geometric mean of the extreme values of the distribution; thus, it combines information on both the minimal and maximal values of the distribution.

The pdf was fitted using Bayesian information criteria. Not only the parameter values were fitted but also the specific pdf was selected in an unbiased way from a large number of known probability mass and density functions according to the Bayesian information criteria. Subsequently, the antimode, the minimum value between two modes of the pdf, were determined whenever a mixture distribution, the sum of two or more probability functions, was selected (see Materials and Methods). The antimode was then used as the threshold to dichotomize the cell population.

The thresholds calculated for a Pcdh gene (Pcdhac2) in the somatosensory neuron dataset differed up to around ten times (Figure 1A). The corresponding ON cells frequencies differed less since few cells have count numbers between the ON and OFF states (Figure 1B). Indeed, all methods correlated highly between each other, even the smallest correlation was high (0.79). The dichotomization with the antimode correlated the most with that using FM or GTME (Spearman rank correlation = 0.85 and 0.86), followed by the VRS (0.79). Similar correlations were obtained for the dopaminergic neuron dataset (Figure S2). Therefore, we applied the GTME to all datasets with TPM/FPKM units. It is important to note that GTME thresholds were calculated also for genes with high bimodality index that yielded unimodal probability density functions, which is often the case, when there are few cells in the OFF or ON expression states (Figure S3).

**Figure 1.**
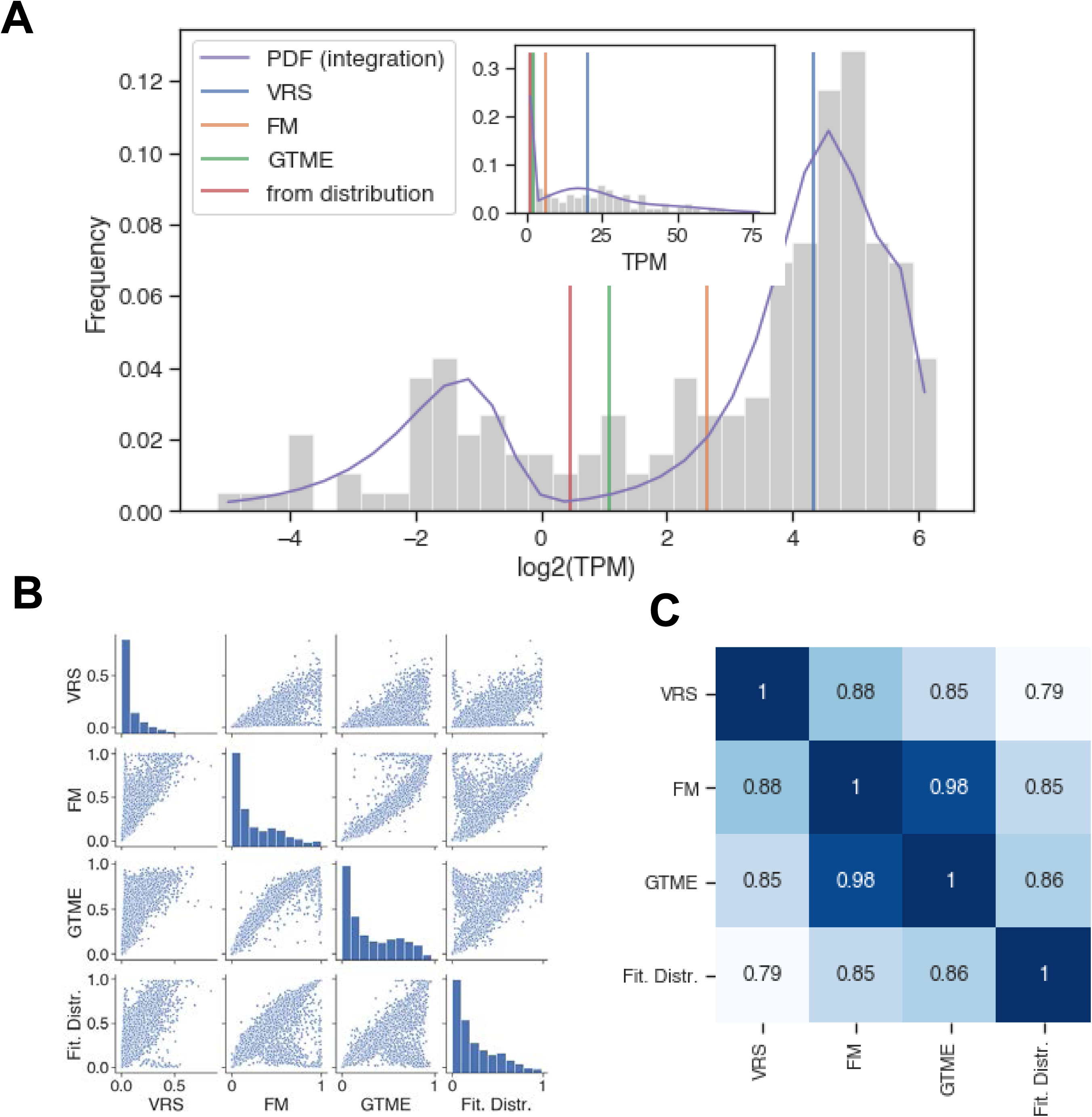
Comparison of dichotomization methods. (**A**) The histogram of the Pcdhac2 transcript numbers in somatosensory neuron. The dichotomization yielded the following thresholds: 2.11 (GTME), 6.28 (FM) and 20.21 (VRS) TPM. The fitted probability density function (pdf) is a mixture of normal distributions, with an antimode at 1.37. The pdf is integrated piecewise according to the logarithmic bins. The inset is a version of the main plot with a linearly scaled x-axis. (**B**) Pair-wise scatter plots showing the ON cell frequencies of each gene in somatosensory neuron dataset with a bimodality index greater than 0.55, after dichotomization with different methods. (**C**) The Spearman rank correlation of the ON cell frequencies shown in (B).

Some datasets had UMI units (Table 1). For these distributions, the Bayesian fitting typically returned Poisson or Yule-Simon distributions, and rarely mixture distributions, which precluded a comparison to antimodes. Therefore, we compared the thresholds according to their ability to dichotomize RNA counts of marker genes of specific cell types (Figure S4). This led to the selection of the FM-threshold for the two datasets. For most genes, the threshold was positioned between zero and one, simply equating the zero expression with the OFF state.

### Effect of chromosomal proximity on stochastic exclusive and concurrent gene choice

Using the dichotomization described earlier, we explored how chromosomal proximity affects interdependence in stochastic gene choice. Chromosomal proximity can influence gene expression in many ways, by promoting the interaction of genes with enhancers via looping, by modifying epigenetic signatures, by relocating chromosomes into active or inactive nuclear compartments, such as transcription factories and heterochromatic compartments (2). These mechanisms may affect only a single gene in a segment but also multiple ones simultaneously. Many of the above mechanisms play a role in the exclusive expression of the Pcdh cluster, the odorant and T-cell receptor genes (6, 34).

If a chromosomal segment alternates between sufficiently large active and inactive nuclear compartments, all or none of the genes in that segment will be expressed, which will result in a large cell-to-cell variation in the number of expressed genes in that segment (Figure 2A, co-occurrence, a.k.a. concurrence). Conversely, the number of the expressed genes can be uniform in a cell population even though each gene is chosen randomly in a cell. While exclusivity is often equated with the expression of a single gene isoform, this is not necessary as long as the overlap among the chosen genes is small (Figure 2A, exclusivity). It is the constant proportion what matters, which is particularly important for protein complexes with fixed stoichiometry such as the Pcdh proteins (10), combining precision with diversity. Alternative chromosomal configurations in which some genes are positioned into active nuclear compartments while the others are prevented from being activated can generate such stochastic exclusive expression.

**Figure 2.**
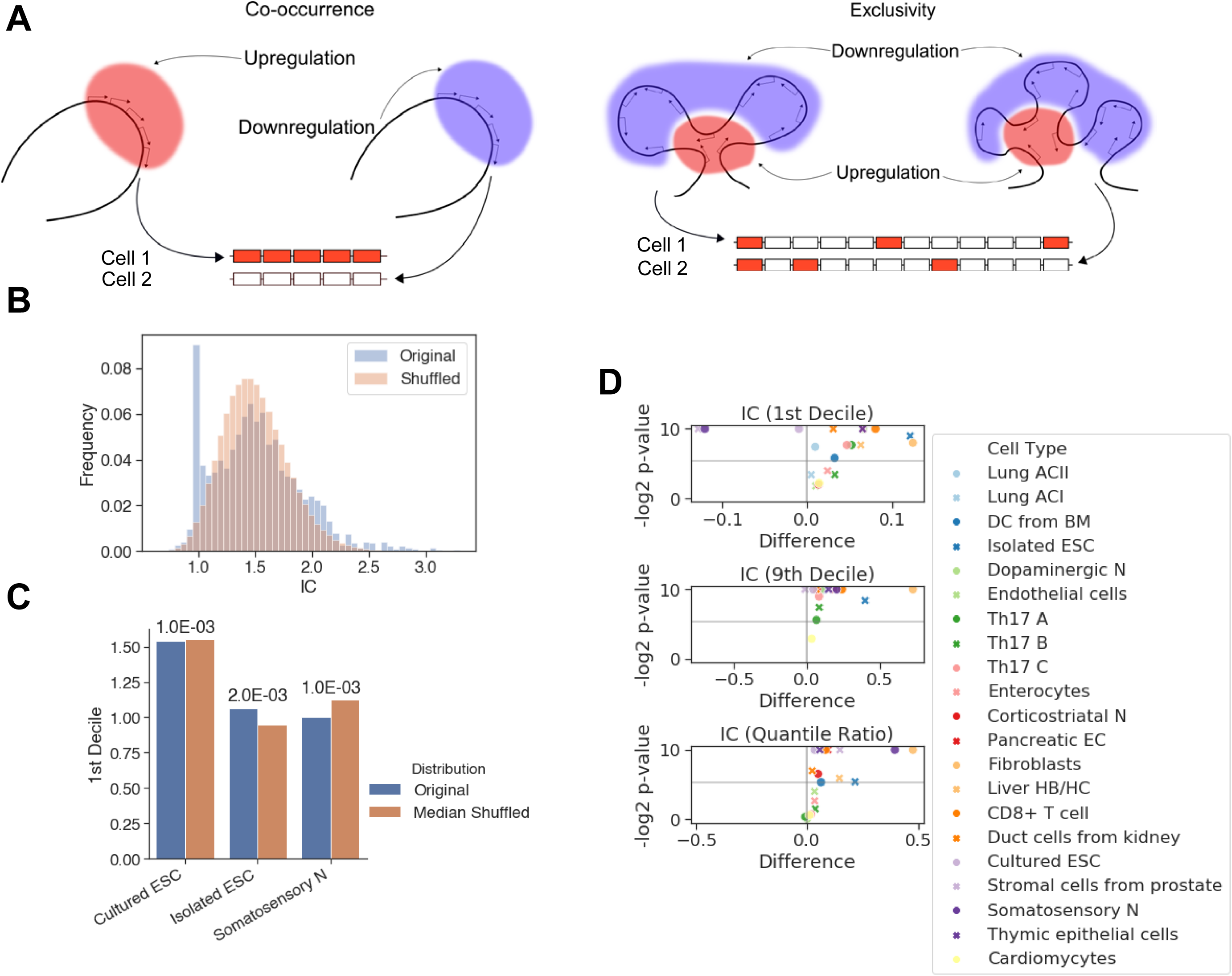
The effect of chromosomal adjacency on stochastic gene choice. (**A**) Schemes showing how the two major forms of stochastic interdependence, the concurrent and exclusive stochastic gene choice, can arise from alternative chromosomal configurations (the variance in the number of expressed genes is 12.5 and 0, respectively). (**B**) The IC distributions calculated from the original and the shuffled genomes with a 14-gene segmentation. There is a significant difference between the two distributions (Kolmogorov-Smirnov test, P-value = 6.22 ·10^−20^). (**C**) The 1^st^ decile IC values are shown for the original and the shuffled genomes, along with the P-values (permutation test) for three cell types. (**D**) Volcano plots showing the difference of 1^st^ Decile, 9^th^ Decile and Quantile Ratio IC values between the original and the shuffled genomes, along with the corresponding P-values (permutation test), performed on the chromosomal segments encompassing 14 genes. The gray horizontal line at 0.025 corresponds to a two-tailed significance level of 0.05.

The stochastic interdependence of the genes under consideration can be quantified with the interdependence coefficient (IC). IC is the ratio of the cell-to-cell variance in the number of expressed genes to the variance of the Poisson-binomial distribution expected from the ON state frequencies of the genes under consideration (see Materials and Methods). An IC less than one indicates exclusivity, while an IC greater than one indicates concurrence. Thus, IC enables the detection of exclusive gene expression even when the mean number of expressed genes is greater than one (as in Figure 2A).

We hypothesized that alternative states of the chromosomes would lead to the overrepresentation of gene segments with exclusivity and concurrence. Therefore, we segmented the chromosomes and calculated the IC for segments comprising 7, 14 or 21 genes sampled along the chromosomes. To test our hypothesis, we reshuffled the genes in the genome and calculated the IC for the segments in the reshuffled genome. By repeating the reshuffling, we obtained a representative distribution of IC values. The distribution of the IC values calculated for the genomic and reshuffled segments differed significantly (Figure 2B). To characterize the differences in the distributions, we compared the location of 10^th^ and 90^th^ percentiles, and their ratio, to assess the enrichment of the exclusive and concurrent segments.

For the somatosensory neurons and stromal cells from the prostate, the 10^th^ percentile IC was smaller for the genomic segments than for the reshuffled segments, which indicates that exclusivity is promoted in these cells (Figure 2C, D). However, exclusivity is suppressed in most cell types (Figures 2D). Interestingly, all cell types, including the somatosensory neurons, were enriched in concurrent segments (Figure 2D). Most genomes display also a significant broadening in the range of IC values, which is mostly due to the overrepresentation of concurrence (Figure 2D).

### Gene segments with stochastic exclusive choice

To characterize the location of the segments that conserve stochastic exclusive expression in multiple cell types, we selected all segments that belong to the bottom 2.5 percentile of the IC distribution in at least two different cell types (or cells cultured in different conditions). These two criteria were extended by a third one, specifying that the IC has to be significantly less than one (i.e. the 95% confidence interval has to be below one) at least in one of the cell types.

Interestingly, the segments overlapping with the Pcdh array represented the largest fraction (Figure 3A, B). Segments from the Pcdh array were identified in all analyzed types of neurons (corticostriatal, dopaminergic and somatosensory), and even in non-neuronal cells, such as the lung alveolar cells. The Pcdh beta isoforms play a role in tumor suppression in lung cancer (35), implying the possibility that exclusive Pcdh expression may diversify cellular identity in non-neuronal cells, as well.

**Figure 3.**
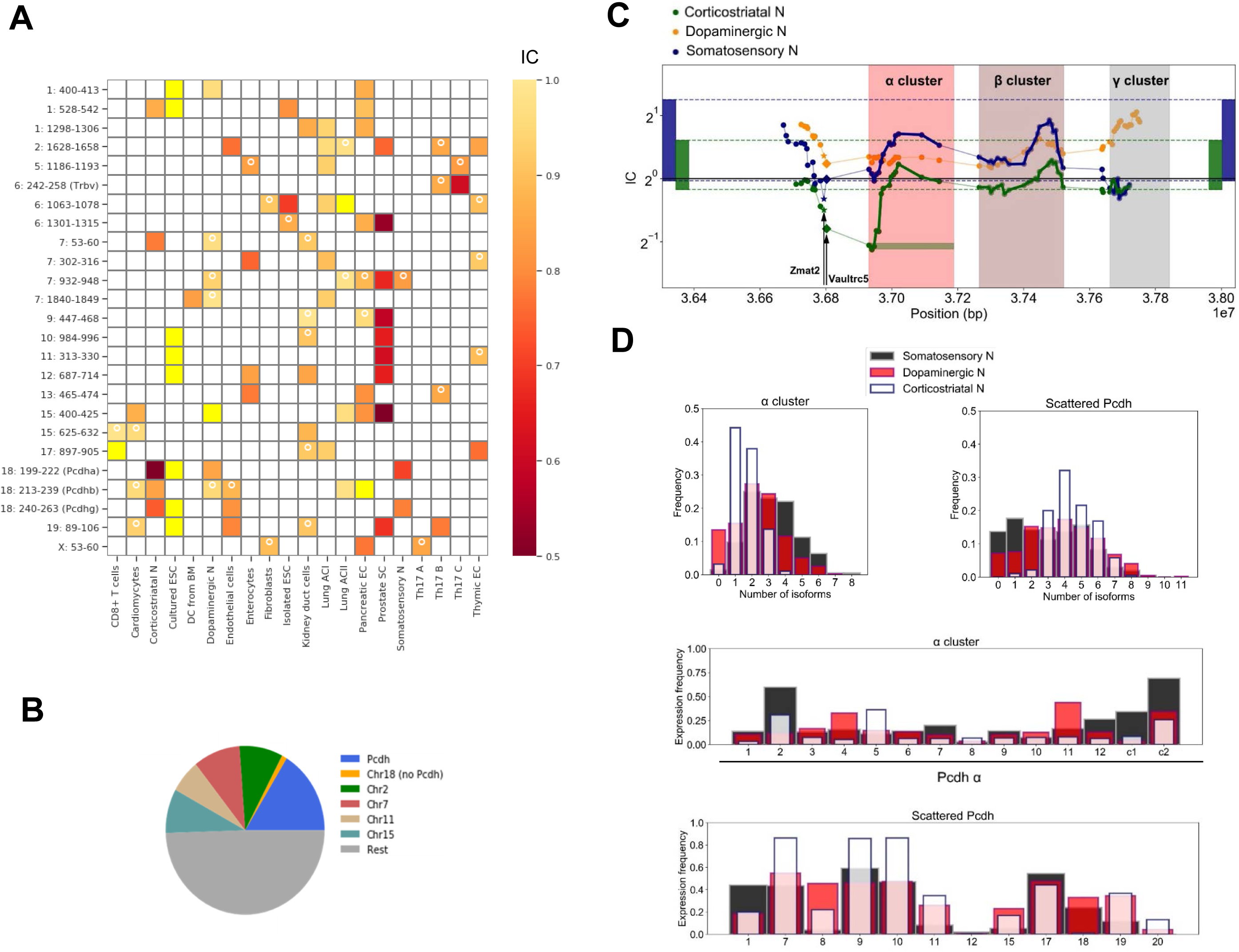
Chromosomal segments with stochastic exclusive gene choice in multiple cell types. (**A**) Chromosomal segments with IC values in the bottom 2.5 percentile of the IC distributions in more than two cell types. Segments having an IC value significantly less than one at least in one cell type are shown provided they express at least 0.03 genes per cell on average and at least 5 gene isoforms in the population of a particular cell type. The white circle denotes segments with an IC numerically less than 1 without reaching significance. In this case, 6 gene isoforms had to be expressed in a cell type. The yellow squares denote IC values greater than one. (**B**) The number of chromosomal segments with exclusive gene choice (as shown in A) in each chromosome for all cell types combined. (**C**) Each dot shows the IC of a segment of 14 genes at the starting (upstream) position. A full segment is denoted by the green horizontal rectangle, at the first gene of the Pcdh α cluster. The two genes upstream of the Pcdh α cluster (Vaultrc5 and Zmat2) are marked with a star and diamond. The rectangles located at the two extremes of the plot indicate the 2.5 and 97.5 percentiles of the IC distribution calculated for the chromosomal segments in the genome. (**D**) The number of expressed genes and the expression frequency of gene isoforms is shown for the Pcdh α-array and the scattered Pcdhs in different neuronal types. The following IC values were obtained for the α-array and scattered Pcdh: 1.11 and 2.80 for somatosensory, 1.27 and 2.06 for dopaminergic and 0.48 and 0.97 for corticostriatal neurons.

The α-array contained the segments with the lowest IC values. Interestingly, the exclusivity extends upstream of the Pcdh cluster involving the Zmat2 and Vaultrc5 genes (Figure 3C), which suggests that they may be also linked mechanistically and/or functionally to the array.

The I.C. values display similar profiles along the chromosome encompassing the upstream region and the α- and β-arrays: IC(corticostriatal) correspond roughly to IC(somatosensory) - 0.5. It has to be noted that the overall IC distribution is shifted toward higher values in somatosensory cells, as indicated by the range delimited by the 2.5 and 97.5 percentiles of the genomic IC distribution. The difference persists even in the reshuffled distribution, which may indicate that at least some of the differences in the absolute values may originate in a systemic intrinsic or extrinsic variable. For example, the procedure used for the isolation of cells and RNA and for the RNA detection can introduce positive correlations extrinsically, which makes the average genomic IC appear larger.

Besides the adjacent α-, β- and γ-arrays, there are Pcdh genes scattered throughout the genome. Thus, the effect of chromosomal adjacency can be specifically assessed for the Pcdh family. The α-array can be conveniently compared with the scattered Pcdhs because they have a similar number of isoforms, 14 and 12, and the mean number of expressed isoforms is similar (3.2 and 3.0 genes in somatosensory neurons, respectively). In both the somatosensory and corticostriatal neurons, α-array belongs to the lowest decile of I.C. distribution, whereas the scattered protocadherins have at least twice as large IC values (Figure 3D).

The chromosome 6 harbors a second prominent gene array, the Trbv, which encodes the variable domains of the T-cell receptor. The low IC values of the overlapping segments indicate a strong exclusivity: it is significantly below one in one of Th17 cell variant and numerically less than one in another Th17 cell variant (Figure 3A). It is important to note that the list of identified arrays with exclusive gene expression is unlikely to be exhaustive because some genes are not detected in a particular cell type. For example, the RNA-seq data cover the expression of Trbv in Th17 cells but not in CD8+ lymphocytes, even though stochastic gene choice and allelic exclusion have been primarily studied in CD8+ lymphocytes. The importance of the exclusivity in T-cell receptor expression in Th17 lymphocytes is underscored by the presence of IL-17 in the cytokine storms, which are thought to contribute to the lethality of the coronavirus disease Covid-19 (36). Dual reactive lymphocytes that recognize endogenous, neurologically relevant, antigens as well as the coronavirus have also been detected (37).

### Stochastic interdependence in gene families

The detection of Trbv and Pcdh arrays based on their low IC values indicates that gene sets with exclusive gene expression can be identified solely based on RNA-seq counts without any information on the alleles and sequence similarity. These gene families have two characteristic features: they are encoded by similar sequences and form an array along the chromosome. The family aspect may be more important for the odorant receptors since the more than a thousand receptor isoforms are encoded by multiple arrays scattered over a large number of chromosomes. Therefore, after having explored the effect of chromosomal proximity, we turned our attention to gene families.

In a study exploring how the expression of odorant receptors changes during cell differentiation, a familywise threshold was used to dichotomize the expression states in order to take into account the lower RNA counts of some isoforms (38). Here, we have adapted the GTME to gene families by combining the information on all the genes in a family to calculate the familywise threshold (see Materials and Methods). This threshold resulted in an IC=0.48 and the mean number of expressed genes was 1.0 (Figure 4A), evidencing a marked exclusivity.

**Figure 4.**
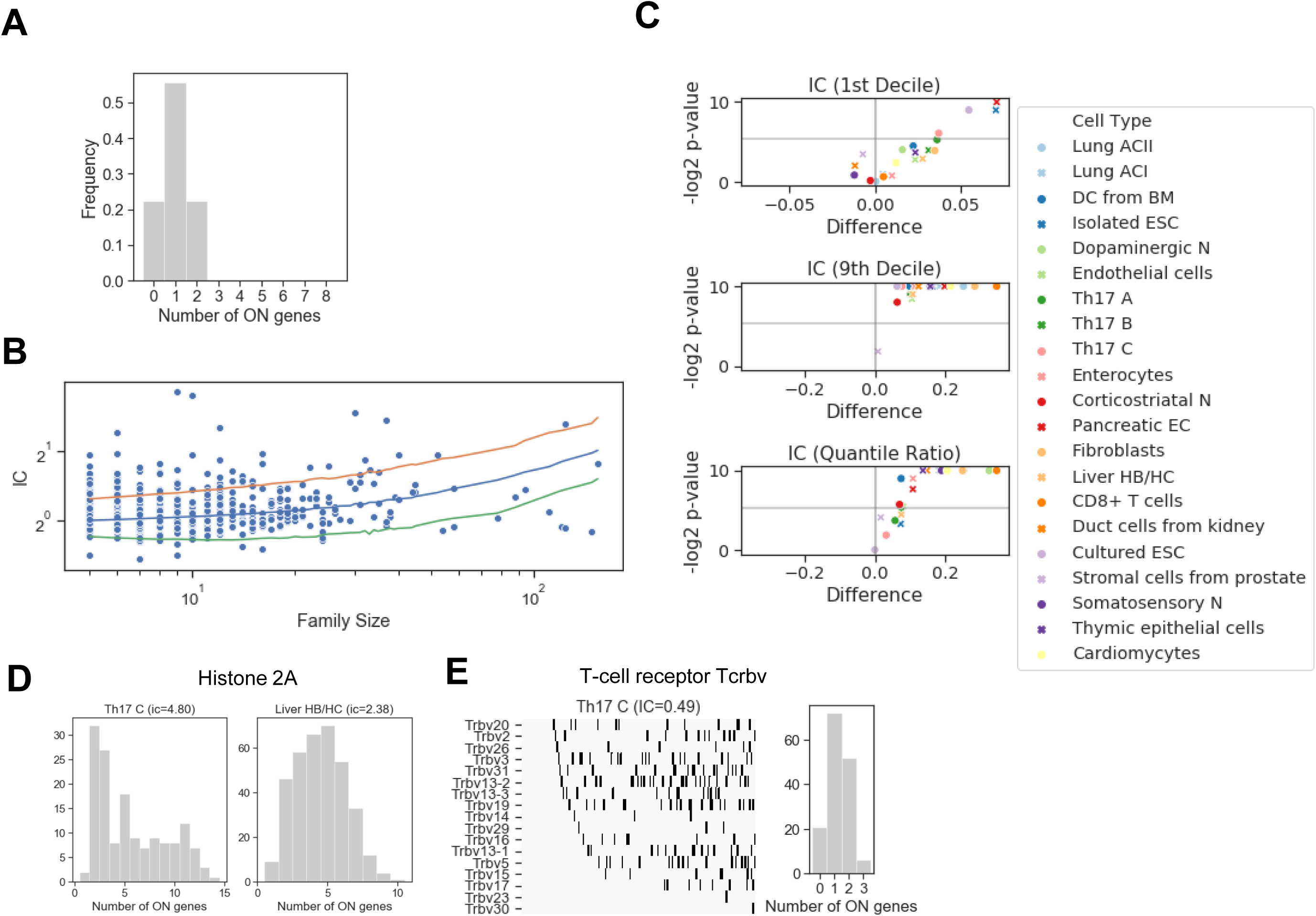
Stochastic interdependence in gene families. (**A**) The number of expressed genes per cell in the family of odorant receptor genes, dichotomized with the familywise threshold (70.7 TPM). Number of cells is *N* = 27. (**B**) IC values of individual families in somatosensory neuron dataset, grouped by the family size. The majority of the families with ICs exceeding either 2.5 or 97.5 IC percentiles of the shuffled genome (orange and green lines, respectively) are concurrent. (**C**) Volcano plots showing the difference of 1st Decile, 9th Decile and Quantile Ratio IC values between the original and the shuffled genomes, along with the corresponding P-values (permutation test) calculated for the gene families consisting of 7 genes. (**D**) The histone 2A family shows co-occurrence in Th17 and liver HB/HC cells, with an IC value of 4.8 and 2.38, respectively. (**E**) The T-cell receptor beta chain family shows a clear exclusivity in Th17 cells (IC = 0.49). The left plot shows the dichotomized expression states.

Subsequently, we dichotomized the expression in each family in the various cell types and reshuffled all the genes belonging to a family encompassing at least five genes (Figure 4B). In the somatosensory neurons, there were many gene families with an IC larger than the 97.5 percentile of the IC distribution the reshuffled genome, but only a few with an IC less than the 2.5 percentile, suggesting that concurrence dominates also in families. Indeed, the systematic examination revealed that the IC at the 10^th^ percentile displayed a significant change in four cell types and the exclusivity was suppressed in each of them. On the other hand, concurrence was promoted significantly in all but one cell type (Figure 4C). Thus, the familywise organization of genes promotes concurrence even stronger than the chromosomal proximity.

### Allelic exclusion and exclusivity in stochastic gene choice

Allelic exclusion is a specific form of exclusivity. It plays a major compensatory role in the expression of sex chromosomes. In order to compensate the double dosage of the X chromosomes in female mice, one of the X chromosomes is inactivated randomly in each cell. Consequently, only one of the gene alleles, the maternal or paternal, is expressed in each cell (39); however, this allelic exclusion is not associated with exclusive gene choice because all genes (gene isoforms) are expressed at one of the chromosomes (Figure 5A). Allelic exclusion occurs after the stochastic promoter choice of the T-cell gene isoform (40). In the protocadherin array, the genes can be expressed monoallelically or biallelically (41). In summary, allelic exclusion may or may not accompany stochastic gene choice in general.

**Figure 5.**
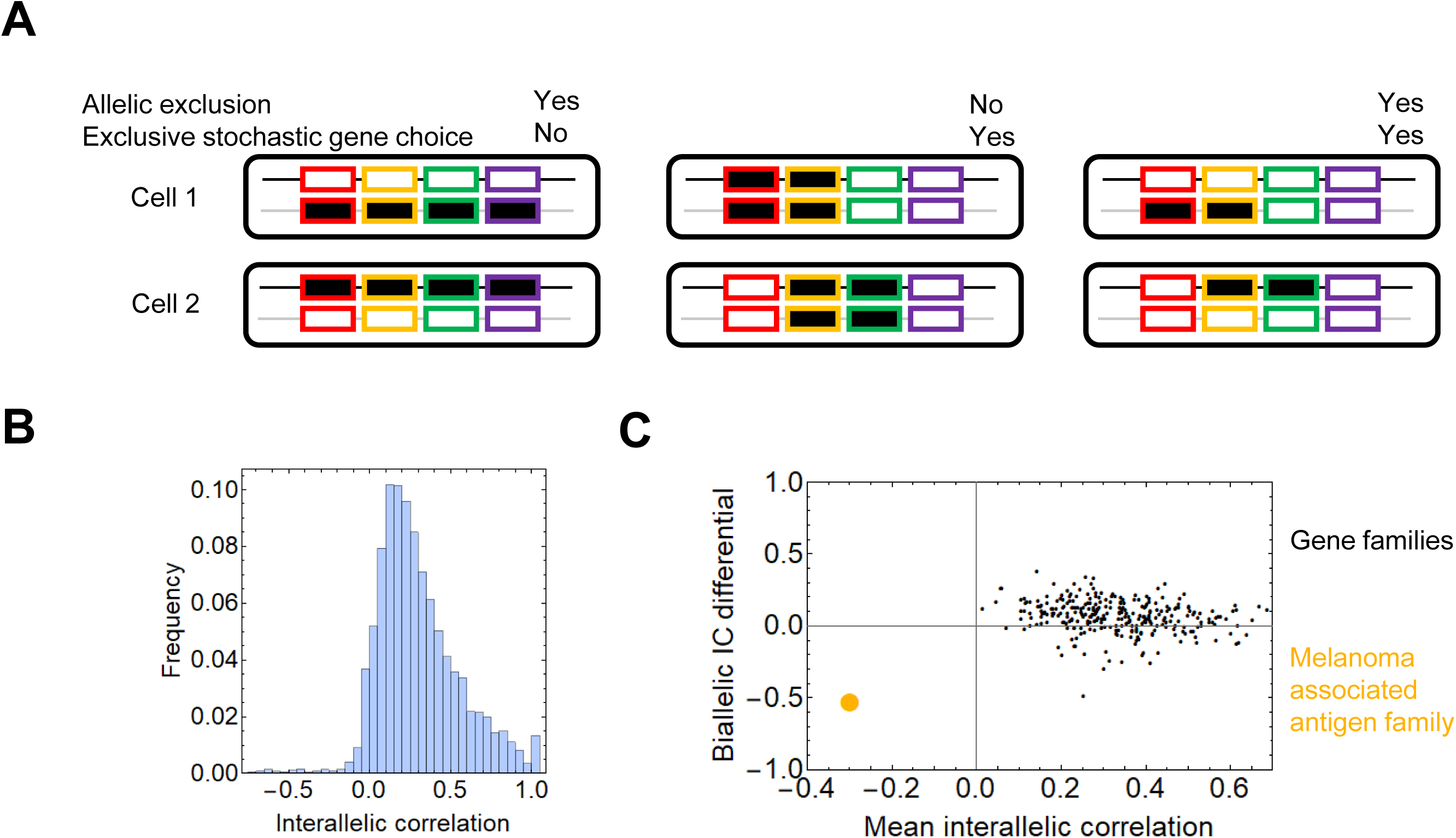
Allelic exclusion and exclusivity in stochastic gene choice. (**A**) Schematic representation of different combinations of exclusivity in allelic and gene choice in an array of four genes. The black and gray lines represent the maternal and paternal chromosomes. The rectangles with no or black filling represent the OFF and ON expression states, respectively (**B**) The interallelic correlation in fibroblasts (42). Negative correlations indicate the allelic exclusion. (**C**) The melanoma associate gene family is highlighted in orange among the gene families. It is the only family with negative mean interallelic correlation.

To assess whether allelic exclusion can contribute to stochastic gene choice, we analyzed RNA-seq data obtained from heterozygous fibroblasts (42), in which the two alleles of most genes can be distinguished. As a measure for allelic exclusion, we calculated the Spearman correlation coefficient between the two alleles for each gene. The overwhelming majority of the genes displayed positive interallelic correlation. Only a small proportion of genes had negative correlation, most of them are located on the X-chromosome, confirming the predominance of this classical form of allelic exclusion (Figure 5B). 17 genes from the β- and γ-arrays of the protocadherin cluster are also expressed; all of them have a positive interallelic correlation with a mean value of 0.53 (Figure S5). Next, we calculated two variables for each gene family: the mean value of the interallelic correlations and the biallelic IC differential (see Materials and Methods). The biallelic IC differential is negative if the IC is reduced upon combining the alleles from the two haplotypes, implying that allelic exclusion contributes to exclusivity in stochastic gene choice in the family. Nearly all gene families have mean positive mean inter-allelic correlation and it does not have a discernible relation with the biallelic IC differential (Figure 5C). The only gene family with negative mean inter-allelic correlation is the melanoma associated antigen family. Interestingly, this family experiences the largest shift toward exclusivity in the stochastic gene choice when the two haplotypes are combined: IC= 2.47 and 2.67 for the haplotypes and IC = 1.52 for the diplotype. Most genes of the melanoma associated antigen family are located on the X-chromosome, and the rest of them at the Prader-Willi locus, which is also known to be imprinted (43, 44), and explains the marked allelic exclusion in this family. These findings indicate that gene families with allelic exclusion are rare; however, specific gene families can utilize it to enhance exclusivity in stochastic gene choice.

### Gene families with stochastic exclusive gene choice

Our comprehensive analysis shows that T-cell receptor beta-chain family in the Th17 cells was the most exclusive among all families, with IC ranging between 0.49 and 0.62 (Figure 4E); similar values are found for the odorant receptor (Figure 4A). The Pcdh family is exclusive in corticostriatal neurons and endothelial cells, in agreement with the findings that only a part of the cluster (typically the α-array) is exclusive and not the entire clustered Pcdh family (Figure 3).

For comparison, the histone 2A family is one of the families with the largest IC values (IC=4.80 in Th17 and 2.38 in liver cells), which nicely illustrates the functional relevance of concurrence. The differentiation state plays also an important role in the expression pattern. Furthermore, some cells enter the S-phase of the cell-cycle and express the histones to support the ongoing DNA replication, while the cells in the other phases of the cell cycle do not express and / or are degraded (45), which results in a large coherent cell-to-cell variation in the number of expressed gene isoforms (Figure 4D).

Thus, our analysis with a simple metric confirmed the exclusivity of all three prototypic families and gene arrays (T-cell receptor, odorant receptor, Pcdh), so they serve as the positive control for the identification of gene families that have exclusive gene expression in multiple cell types. We used the same criteria as for the chromosomal segments.

The majority of the retrieved families encode membrane proteins (Figure 6A, 7A) like the three prototypic families. Many of them are associated, directly or indirectly, with two processes: transmembrane ion transport and intercellular adhesion (Figure 6A), exemplified by the sodium/potassium transporting ATPase subunit gamma, the basigin related families, and jointly the carbonic anhydrase and anion exchange proteins (Figure 7B).

**Figure 6.**
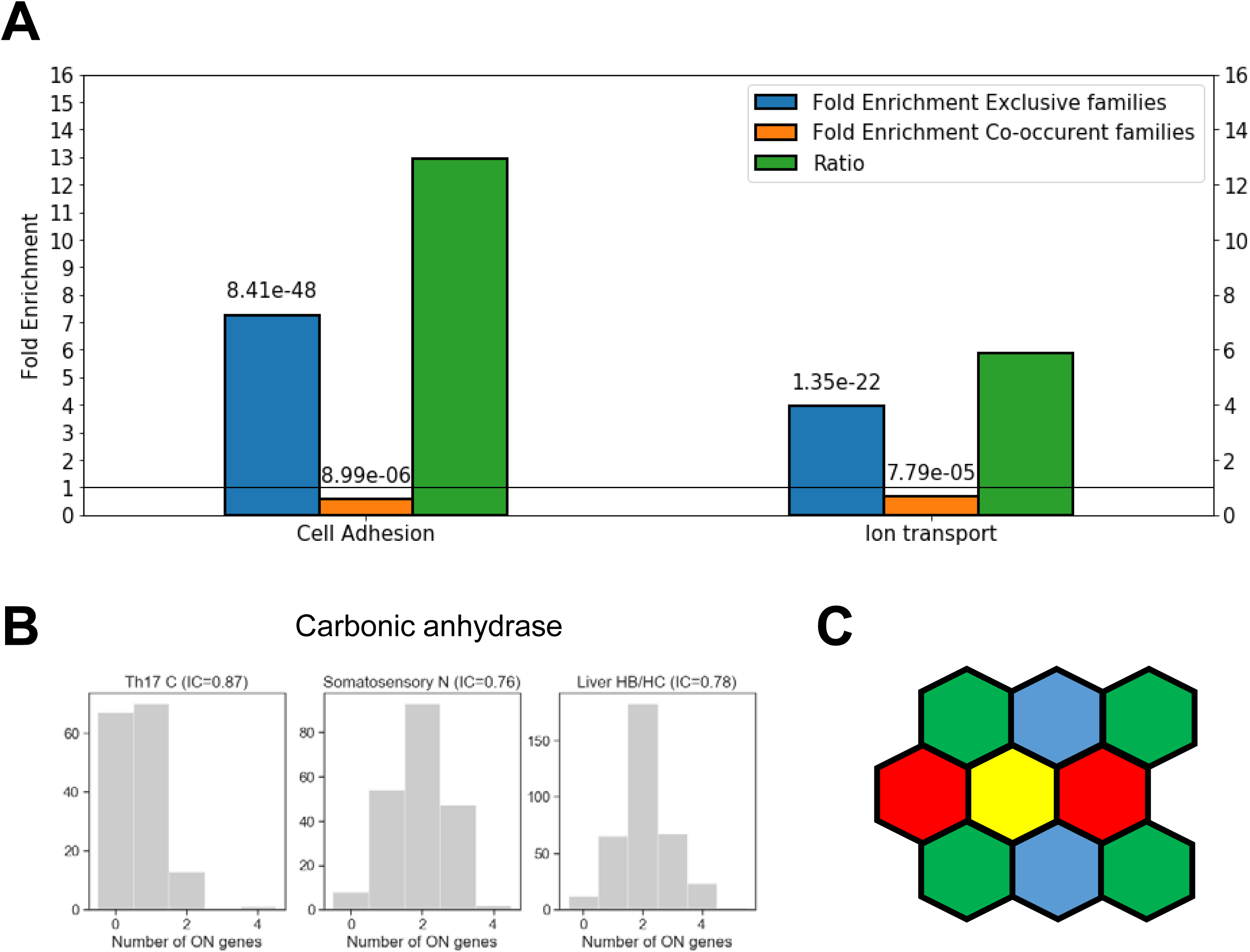
Cellular individuality and cell adhesion. (**A**) Enrichment analysis for genes belonging to exclusive and concurrent gene families. The P-values are indicated on the top of the bars. The exclusive families were selected with the same criteria described in Figure 4B. The concurrent families (IC belonging to top 2.5 percentile) were constrained further with the following criteria: mean number of expressed genes per cell higher than 0.03, IC significantly higher than 1 and at least 5 non-zero genes per family. We considered all the genes expressed at least in one cell type belonging to the selected families. The enrichment analysis was performed though an enrichment analysis tool (http://geneontology.org/). The figure shows two selected functions: Cell adhesion (GO: 0007155) and Ion transport (GO: 0006811). The ratio of the fold-enrichment in the exclusive to that in concurrent families is shown. (**B**) The number of expressed carbonic anhydrase genes per cell in Th17 cells, somatosensory neurons, and liver HB/HC (IC = 0.87, 0.76 and 0.78, respectively). (**C**) Schematic representation of cells in a place showing that the exclusive expression of four gene isoforms (colors) is sufficient to confer cellular individuality.

**Figure 7.**
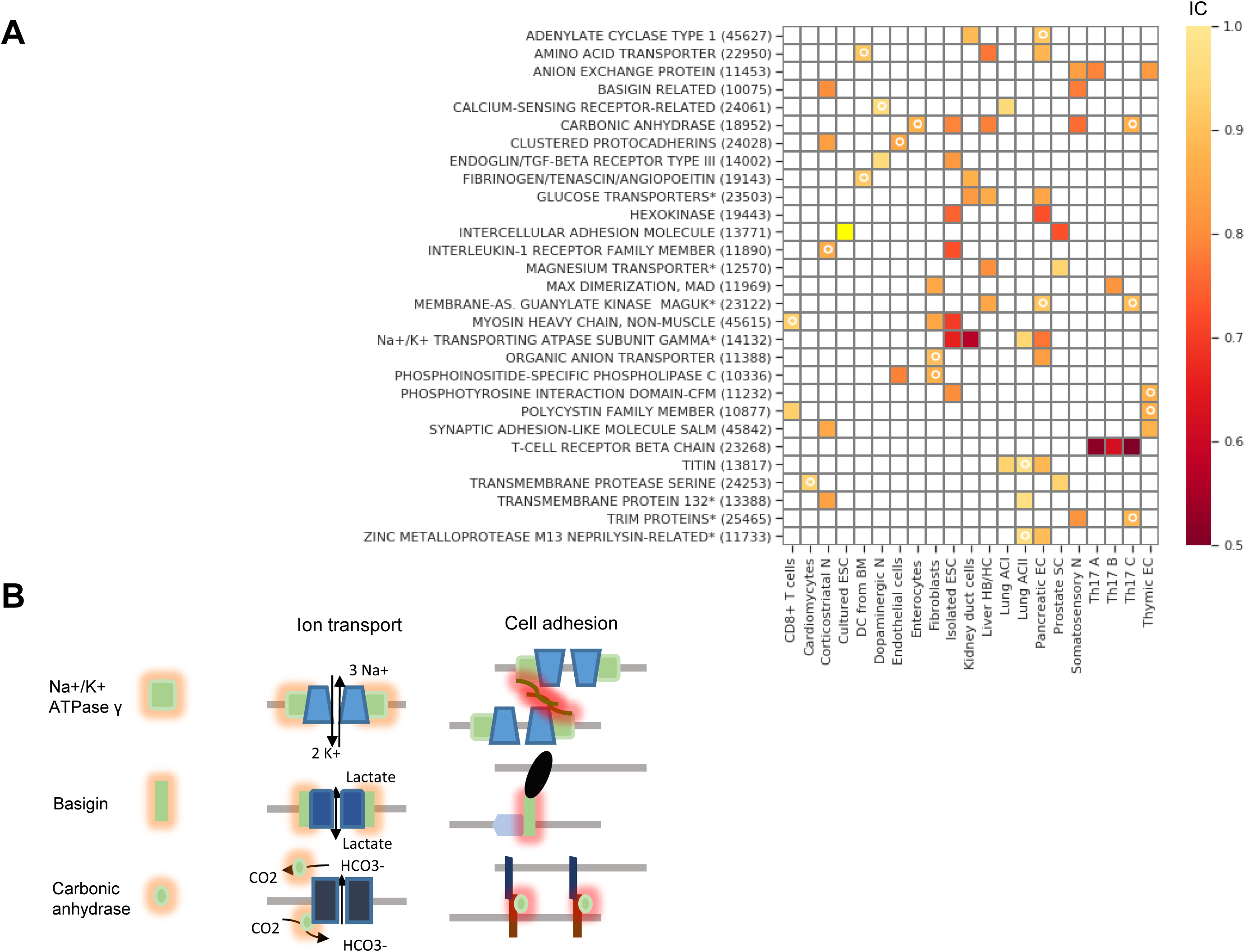
Gene families with exclusive gene expression. (**A**) Gene families with stochastic exclusive gene choice in two or more cell types; further details of selection as in Figure 3 (see also Data S1). For the families labeled with star, descriptive names were given instead of the Panther names. The panther numbers of the families are indicated in parenthesis. (**B**) Schematic representation highlighting the dual role of three gene families (Fxyd, basigin and carbonic anhydrase genes). On the left side, the cis interaction of the corresponding proteins with channels and pumps is denoted by orange shades. These functions are related to metabolic and ion homeostasis. On the right side, the trans-interaction with ligands on the adjacent cells is labeled with red shades. The glycosylation of the Fxyd protein affects the transdimerization of the Na^+^/K^+^ ATPase. The carbonic anhydrase interacts with the anion exchange protein, which transports HCO_3_^-^.

The Fxyd1-7 gene isoforms encode the gamma subunit of the Na^+^/K^+^ ATPase, which is the regulatory subunit of this ion pump. While these ATPases are primarily involved in ion homeostasis, they can also trans-dimerize and thus mediate cell-to-cell interaction. It has been shown that the ratio of the Fxyd5 isoform to the α1–β1 heterodimer determines whether the Na^+^/K^+^ ATPase acts as a positive or negative regulator of intercellular adhesion (46). This is highly reminiscent of the Pcdh proteins, in which the ratio of the expressed isoforms determined intercellular adhesion (10, 47). It is important to note that cells in a plane can attain complete cellular individuality with the exclusive expression of four different gene isoforms, according to the four color theorem (Figure 6C) (48). Somewhat higher numbers are needed for cells arranged in 3-dimensional interaction networks.

Interestingly, basigin can also bind the β2-subunit of Na^+^/K^+^ ATPase (49). The stochastic exclusivity of the basigin-related genes was detected in somatosensory and corticostriatal neurons (Figure 7A). The members of this family are named after the immunoglobulin-superfamily molecule basigin and are well known mediators of intercellular adhesion (50), comprising genes such as Contactin 6 (Cntn6), Down syndrome cell adhesion molecule (Dscam) and Neuronal cell adhesion molecule (Nrcam). The basigins often interact with monocarboxylic acid transporters, which catalyze the transport of lactate, pyruvate, etc. (51); thus, they indirectly affect the ion transport.

The carbonic anhydrase family displays a similar duality of functions related to ion homeostasis and intercellular adhesion, and have a pronounced exclusivity (IC between 0.76 and 0.87, Figure 6B). The primary role of carbonic anhydrases is the catalysis of the reversible conversion of CO_2_ to carbonic acid. However, some isoforms have lost their catalytic activity (Car8, 10 and 11) and they play a role in promoting the diversification in neuronal interactions. The Car10 and Car11 isoforms are secreted glycoproteins that are predominantly expressed in the brain. Car10 was shown to be a conserved pan-neurexin ligand (52). Neurexins, like protocadherins, mediate interneuronal interactions, but the isoform diversity is generated primarily through alternative splicing (53) and not by stochastic gene choice. Overexpression of Car10 in neurons creates a shift in neurexin isoforms in mouse and human neurons, which may explain how the stochastic choice of Car isoforms generates diversity. Even catalytic Cars affect intercellular adhesion. For example, Car9, a cancer associated transmembrane isoform of carbonic anhydrase, reduces E-cadherin mediated adhesion (54). The Cars can interact with the anion exchange proteins, Slc4a, which transport bicarbonate (55), which is thought to accelerate CO_2_ transport. Thus, two families with exclusive expression can interact physically.

Two other families with exclusive expression interact in a similar way. The glucose transporters and the phosphorylation of glucose by the hexokinases act in two consecutive steps of the glycolysis pathway. The remaining family with marked exclusivity, the non-muscle myosin heavy-chain, is not a membrane protein but may affect cell adhesion (see Discussion).

## DISCUSSION

Exclusive gene choice has been equated with the one gene isoform per cell rule. The increasing number of exceptions to this rule and their physiological relevance necessitate formulating alternative definitions that can be used for multi-subunit proteins, as well. Since the IC is a relative measure it can be used for multi-subunit proteins. An IC less than one implies that the gene choice is more precise than by random, independent selection. With this simple definition, we show that T-cell receptors and odorant receptors belong to the families with the highest degree of precision (IC ∼ 0.5) despite the presence of some cells that evade the one-receptor per cell rule. Furthermore, when the most exclusive segments are combined from all cell types including even non-neuronal cells, the segment containing the Pcdh array is the most prominent. The Pcdh α-array expresses two or more isoforms in most neurons.

Thus, the ranking of families according to the IC retrieves the families according to classical definition. Thus, this correspondence validates the approach relying on the Poisson-binomial distribution.

Our results show that stochastic exclusivity is rare in both gene families and segments and concurrence is overrepresented. Multiple mechanisms are likely to underlie this phenomenon. Evolving from a single gene, paralogs have common regulatory sequences. Consequently, a shift from concurrence toward exclusivity is expected only after a sufficient evolutionary divergence in the family. Chromosomal proximity can also promote concurrence when a transcription factor affects multiple genes in a chromosomal segment (56). For example, two copies of the same gene at the same chromosomal position experience more correlated fluctuations if they are positioned on linked chromosomes than on physically separated, but homologous, chromosomes (57). Furthermore, the positive correlation in stochastic gene expression has gradient-like features along the mammalian chromosomes (58). Thus, the predominance of concurrence in the genome can be viewed as a direct consequence of the evolutionary-genetic and biophysical-chemical processes.

Despite the dominance of concurrence in chromosomal segments, chromosomal proximity may promote exclusivity provided adjacency is placed in the appropriate context. A single gene among the adjacent linked Pcdh α-genes can be chosen exclusively upon the formation of a CTCF-mediated chromosomal loop between the chosen gene and a downstream enhancer (59). In agreement with this observation, the Pcdh α-array displays a much higher exclusivity than the scattered Pcdhs, underscoring the importance of having chromosomal adjacency in the appropriate context. Interestingly, several genes with exclusive expression identified in this study, such as Car2 and Slc4a, which are involved in carbonate production and transport, have been shown to be regulated by CTCF (60, 61).

The quantification of stochastic exclusivity with the IC has multiple advantages. It can help to define the range of chromosomal segments subject to exclusive gene choice, especially when the genes do not belong to a family. For example, the exclusivity in the α-Pcdh array extends beyond the array and affects two upstream genes. Interestingly, Zmat2, one of the two genes, has been shown to regulate the splicing of genes involved in cell adhesion (62). Thus, Zmat2 may directly affect the Pcdh-mediated cell adhesion. Similarly, the IC formalism does not require predefined sets of genes for the assessment of exclusivity. For example, the αC1 and αC2 isoforms are usually excluded from the analysis when the number of expressed gene isoforms is quantified in α-array the due to their constitutive expression in Purkinje cells (11). However, their expression is not constitutive in other cell types: the αC1 and αC2 isoforms are expressed at a lower frequency than some of the variable isoforms (α1-12) in corticostriatal neurons (Figure 3D). Since the IC formalism does not assume a single gene to be expressed in order to be exclusive, it permits the detection of exclusivity in all these cell types with different mean number of expressed genes.

In addition to defining the length and location of chromosomal segments with stochastic exclusive expression, the IC has another important aspect, the absolute value. The T-cell receptor family with IC values as low as 0.5 has an unmatched degree of exclusivity in comparison to the other detected exclusive families. This may reflect the fact that multiple different molecular mechanisms cooperate to stabilize exclusive stochastic gene expression: the promoter choice through chromosomal looping is followed by DNA recombination and allelic exclusion (6). DNA recombination is unlikely to contribute to the exclusivity in the families involved in cell adhesion. Since we analyzed genes, driven by distinct promoters, regulation of transcription is likely to be a major contributor to exclusivity for the families detected in this study. The exact mechanism, looping or covalent epigenetic modifications or other processes, remains to be determined.

We have used a relatively stringent criterion to identify segments or families with exclusive choice since it had to be detected in at least two different cell types. Despite the overrepresentation of concurrence, the exclusive gene choice is not restricted to the T-cell receptor, odorant receptor and Pcdh families. Ten other families were identified with pronounced exclusivity, with IC less than 0.8: the anion-exchange and basigin related proteins, the carbonic anhydrases, intercellular adhesion molecule, interleukin-1 receptor family, phospholipase C, the sodium/potassium transporting ATPase gamma subunit, the hexokinases and the non-muscle myosin heavy-chain. Most of them are directly affect cell adhesion (Figure 7), but even hexokinases can affect motor or cytoskeletal proteins, and thus regulate cellular adhesion (63, 64). Ion transport is the second most overrepresented function in the detected families. Ions have been long known to modulate cell adhesion (65). It remains to be determined, whether the functions coupled to cell adhesion, the ion or metabolite transport, profit from the combination of diversity and precision. Recent advances in the description of the spatial variations in metabolism across a cell-population (66, 67) do suggest that not only cell adhesion but also ion homeostasis may profit from stochastic exclusive gene choice.

The functional consequences of diversity through gene choice can be illustrated with membrane proteins that mediate cell adhesion in a way that the affinity of the interaction depends on the particular combination of the respective protein isoforms (10). For example, choosing two isoforms from a repertoire of five genes permits ten combinations, and thus ten cellular identities. The cells assume their identities according to their distinguishability from their neighbors, which they interact with through homophilic or heterophilic interactions (Figure 6C) (47, 68, 69). This combinatorial diversity due to the random choice of multiple gene isoforms is translated into a diversity of cell-to-cell interactions, while the exclusivity guarantees the precise stoichiometry within the membrane protein complexes. This principle is a conserved property of many gene families involved in cell adhesion beyond the protocadherins, suggesting that stochastic exclusive gene choice is an ideal mechanism to link diversity with precision in cell adhesion.

## MATERIALS AND METHODS

### Data sources

To define the chromosomal segments, the Genome Reference Consortium Mouse Build 38 patch release 6 (GRCm38.p6) was used [https://www.ncbi.nlm.nih.gov/assembly/GCF_000001635.26]. The genes that are marked as predicted were excluded, and only the genes sourced from Best-placed RefSeq (BestRefSeq) and Curated Genomic were considered.

PANTHER15.0 was used to map genes to their corresponding gene families, [ftp://ftp.pantherdb.org/sequence_classifications/current_release/PANTHER_Sequence_Classification_files/PTHR15.0_mouse] (70). The single cell RNA-seq datasets are described in Table S1.

### Interconversion of RNA-seq quantification units

TPM (Transcripts Per Million) units were analyzed without conversion. The RPKM (Reads Per Kilobase Million) and FPKM (Fragments Per Kilobase Million) can differ between samples, causing biases for the statistical interpretation of the data (71). Therefore, they were converted into TPM units (72):

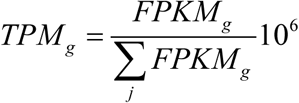

*FPKM*_*g*_ represents the FPKM values of a given gene. The gene counts are summed over the population of *j* cells.

Datasets with Unique Molecular Identifier (UMI) counts were used without further normalization.

### Dichotomization of expression into ON and OFF states for the genes in the chromosomal segments

TPM distributions were filtered to exclude the genes with unimodal expression. For this purpose, the Bimodality Coefficient was calculated for each gene:

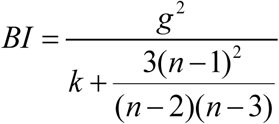

where k is the sample excess kurtosis, g is the sample skewness, n is number of samples (i.e. cells) [SAS Institute, 1990]. Only the genes with BI higher than 0.55 were kept since a value of 5/9 or less corresponds to a unimodal distribution.

Three methods were compared to dichotomize the expression of individual genes: VRS, FM and GTME. The minimum threshold was set to be 0.5 TPM, which is widely used as threshold for a gene considered to be expressed. Thus, when a procedure resulted in a threshold with a value less than 0.5 TPM, it was replaced by 0.5 TPM. Upon determining the threshold, the genes are dichotomized. If the expression value is greater than or equal to a threshold, the gene is marked as expressed in this cell (i.e. with 1), otherwise it is marked as not expressed (i.e. with 0).

### Variance Reduction Score (VRS)

VRS is a measure of bimodality, in that it reflects how much the variance of the original distribution is reduced in comparison to the sum of the variances of the two distributions obtained by the splitting of the original distribution with a threshold (12).

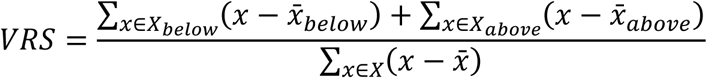

where *X* is a total set of expression values of a gene, *X*_*below*_ and *X*_*above*_ are sets of expression values lower than and greater than or equal to a threshold respectively 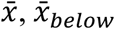 and 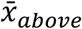 are the mean expression values for the three sets respectively.

In order to find the threshold with the minimal VRS, a range of threshold values were tested for each gene. This range is a list of geometrically progressing series with the step of 1.2 starting at 0.025 quantile of non-zero expression values up to the 0.975 quantile to get a more granular view of VRS at lower thresholds. The threshold that yields the minimum VRS is chosen as a dichotomization threshold.

### Fraction of Maximal values (FM)

The FM is a biochemically motivated threshold and assumes that the expression of a gene does not vary too much around its activity specific to the ON state. For this purpose, the 1/10th of the TPM value at the 97.5 percentile was chosen. If the number of cells with non-zero expression values (*N*) is less than 120, then the 1/10th value of the average (arithmetic mean) of the three largest values was calculated.

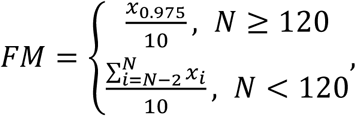

where *x*_*p*_ is the *p*th quantile of non-zero expression values, *x*_*i*_ is the *i*th element of the sorted non-zero expression value list, *N* is the number of non-zero expression values,

### Geometric trimmed mid-extreme (GTME)

The GTME is motivated by the predictions of transition rates in bistable systems: the threshold between the two states is defined as the geometric mean of the low and high states (73). Bistable systems can underlie bimodal distribution but there is no simple relation between them because of the transiency (74). In order to define the threshold without knowing the exact values of ON and OFF states, the geometric mean of the non-zero TPM values at the bottom and top 2.5 percentiles (40-quantiles) of the distribution were taken. If the number of non-zero TPM values is less than 120, the average (arithmetic mean) of the three least and largest values were used to calculate the geometric mean.

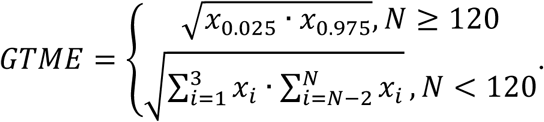

Analogous thresholds allow for the precise calculation of the transition rates in a bistable cell population (73).

### Familywise Thresholds

Assuming that the expression values of genes within a family are similar, a common threshold can be defined for all genes within a family. The familywise FM (fFM) and GTME (fGTME) were calculated as follows. The RNA counts larger than 0.5 were considered instead of the x>0 condition. When the respective cell number *N* was larger than 120, the *x*_g, 0.025_ and *x*_g, 0.975_ were calculated for each gene. The fFM was calculated from the maximum of the set of *x*_g, 0.975,_ *g* ∈ *GF*, representing each gene in a gene family (*GF*). Thus, a single gene in the family determines the threshold for all the genes in the family. Similarly, the two genes corresponding to the minimum of the *x*_g, 0.025_ and the maximum of the *x*_g, 0.975_ *g*∈ *GF*, set determine the fGTME. Analogous calculation were performed for *N*<120, with mean averages of the three largest and smallest expression values, instead of the values at the percentiles.

### Fitting of distributions

Probability density (or mass) functions, *φ(x*), were fitted with the FindDistribution of Wolfram Mathematica, which combines the Bayesian information criterion with priors over distributions to select both the best distribution and the best parameters for it. Commonly fitted distributions were the Binomial, Cauchy, Exponential, Gamma, Geometric, Normal, Laplace, Logistic, Lognormal, Poisson, Negative Binomial, Yule-Simmons distribution and their mixtures. Whenever a mixture distribution was obtained by the FindDistribution, the antimodes were calculated. The antimodes were determined analytically based on the first and second derivatives of *φ*(*x*). The smallest antimode in the range *x* > 0.5 was used as thresholds for dichotomization for each gene. As opposed to other methods, the *φ*(*x*) based thresholds were not used for calculation of IC across the genome, since they were obtained for a smaller number of genes in comparison to the other methods.

### The interdependence coefficient (IC)

The IC is the ratio of the observed variance in the number of expressed genes in a cell population to the variance of the Poisson binomial distribution expected from the expression frequencies (13). The variance of the generalized binomial (Poisson-binomial) distribution is a function of the probability of each isoforms *i* to be expressed (*p*_*i*_):

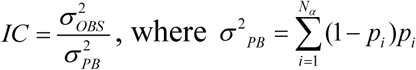

*p*_*i*_ is equal to the ON cell frequency. IC=1 indicates an independent stochastic gene choice according to the Poisson-binomial distribution, akin to a relation Fano-factor=1, which indicates a Poisson distribution for a single gene (75).

The 95% confidence interval (CI) of the IC was calculated by bootstrapping. After resampling the cell population, the observed variance and the expected Poisson-binomial variance were calculated for each resampling, and IC was calculated. When the 95% CI was below one, exclusivity was considered significant.

### Permutation tests

Permutation tests were used to assess the effect of chromosomal adjacency and family membership on stochastic interdependence. The expression values of the genes are shuffled among all genes but for those that were not measured in a particular dataset or were not bimodal. The shuffling was performed 1000 times. Similarly, the assignment of genes (i.e. their respective expression values) to gene families is shuffled. Only the genes that are present in both the families and the RNA-seq datasets are reassigned in a way that the sizes and number of families are preserved. The distribution of IC values were obtained for each re-shuffling.

Firstly, the Kolmogorov-Smirnov (KS) test was performed to see if the original distribution is different from the concatenated shuffled distributions. The implementation of the test provided by SciPy was used.

The P-value of KS test lower than 0.05 indicates the significance level to which the distributions are different; however, the KS test does not show what mode of interdependence is over- or underrepresented. Therefore, the P-values for the changes in the quantiles were calculated based on the permutation tests (76). The 10^th^ and 90^th^ percentiles and their ratio were calculated as representative quantiles for the exclusivity and concurrence. The P-value was calculated as follows

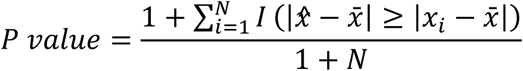

where 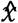 is the original statistic, 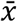 is the mean of the shuffled statistic, *x*_*i*_ is the statistic of the ith permutation, and *N* is the number of permutations. The pseudocount is added to avoid P-values of 0.

A two-tailed P-value of 0.05 was selected for a statistic to be considered significantly higher or lower than the statistic of the shuffled distributions. The original statistic, the median of the statistic for the shuffled distributions and the P-value is reported. Exclusivity is promoted when the 10^th^ percentile of the original distribution is significantly smaller. Similarly, co-occurrence is promoted when the 90^th^ percentile is significantly greater. These tests were applied for each chromosomal segment size separately. Families were grouped according to their size, and the same tests were performed as for the chromosomal adjacency. Only family sizes that have 30 or more gene families were taken for KS and permutation tests.

### Identification of genes subject to concurrent or exclusive gene choice in multiple cell types

To assess which sets of genes conserve their mode of interdependence across multiple cell types, the pairwise overlap of gene segments or gene families that are within the bottom or top 2.5 percentiles of their respective IC distributions was determined. In other words, a segment or a family is considered a hit, if it appears in two datasets in the respective tails of IC distributions. The chromosomal segments were overlapped separately for each segment size, whereas all families were considered together (Data S1). Further conditions to filter the selected genes are described in the relevant context.

### Examination of the relations between allelic exclusion and stochastic gene choice

The mean interallelic correlation was calculated by averaging the Fisher transform of the Spearman correlation coefficient calculated for the two alleles, followed by a back transformation (77):

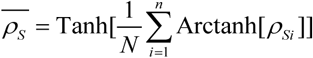

To calculate the biallelic IC differential between the diplotypes and haplotypes, the following formula was used:

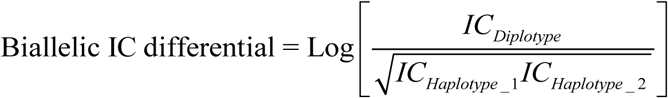

## ACKNOWELDEGEMENTS

We thank Vincent Jaquet, Michael Stadler and Peter Scheiffele for helpful discussions.

## SUPPORTING INFORMATION

**Figure S1.**
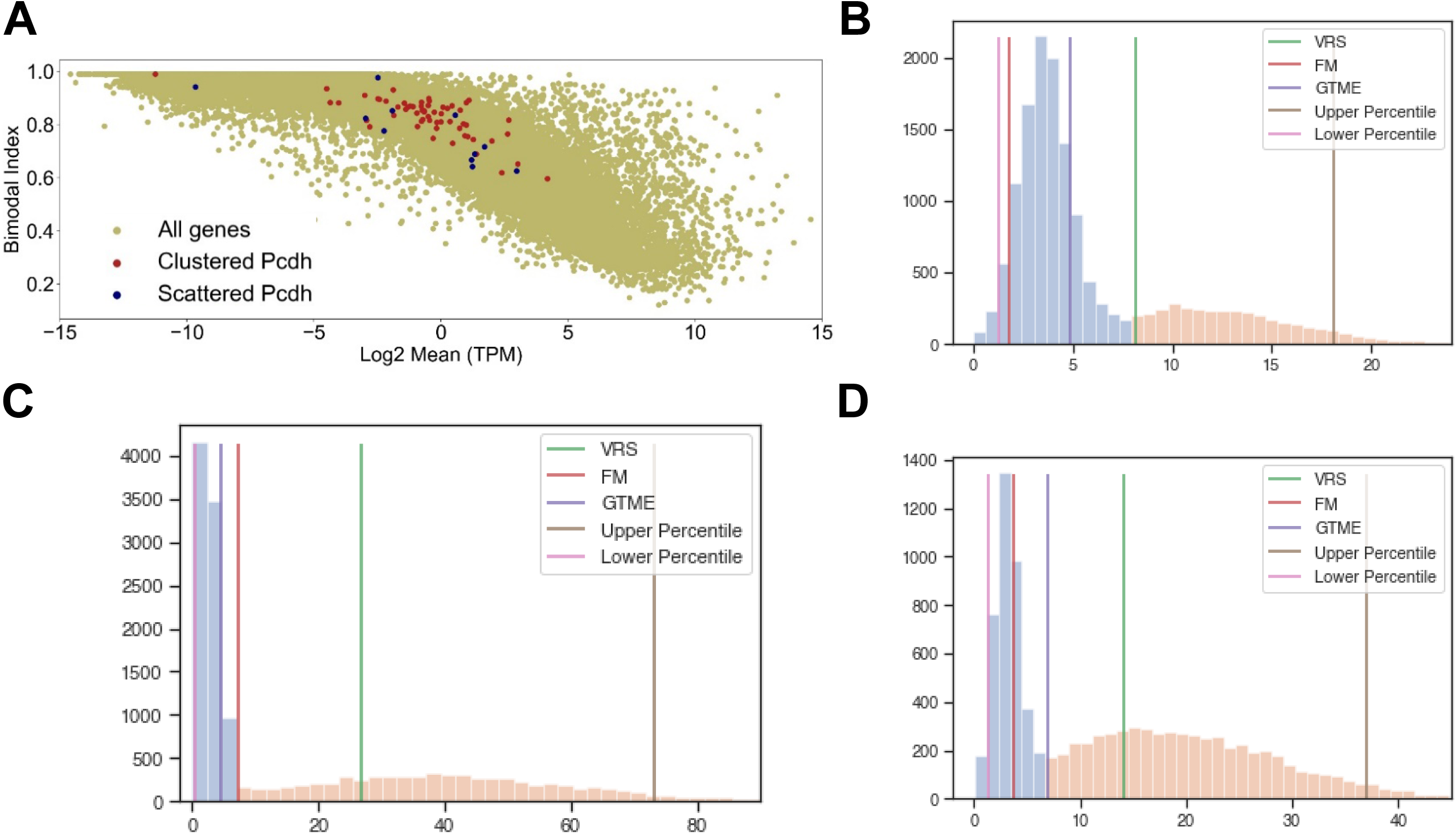
Bimodality and thresholds to dichotomize the distributions. (**A**) The bimodality index calculated for the genes the somatosensory neuron dataset. (**B-D**) Simulated bimodal distributions representing mixtures of two normal distributions *N*(mean, standard deviation). The blue and orange parts denote the distributions left and right to the antimode, respectively. Varying the parameters of the distributions, the best threshold is obtained by different dichotomization methods. (B) *N*(3.5, 1.2) and *N*(11, 5) with a 2:1 proportion. (C) *N*(2, 2) and *N*(38, 22); 10:7. (D) *N*(3, 1) and *N*(17, 11); 3:7.

**Figure S2.**
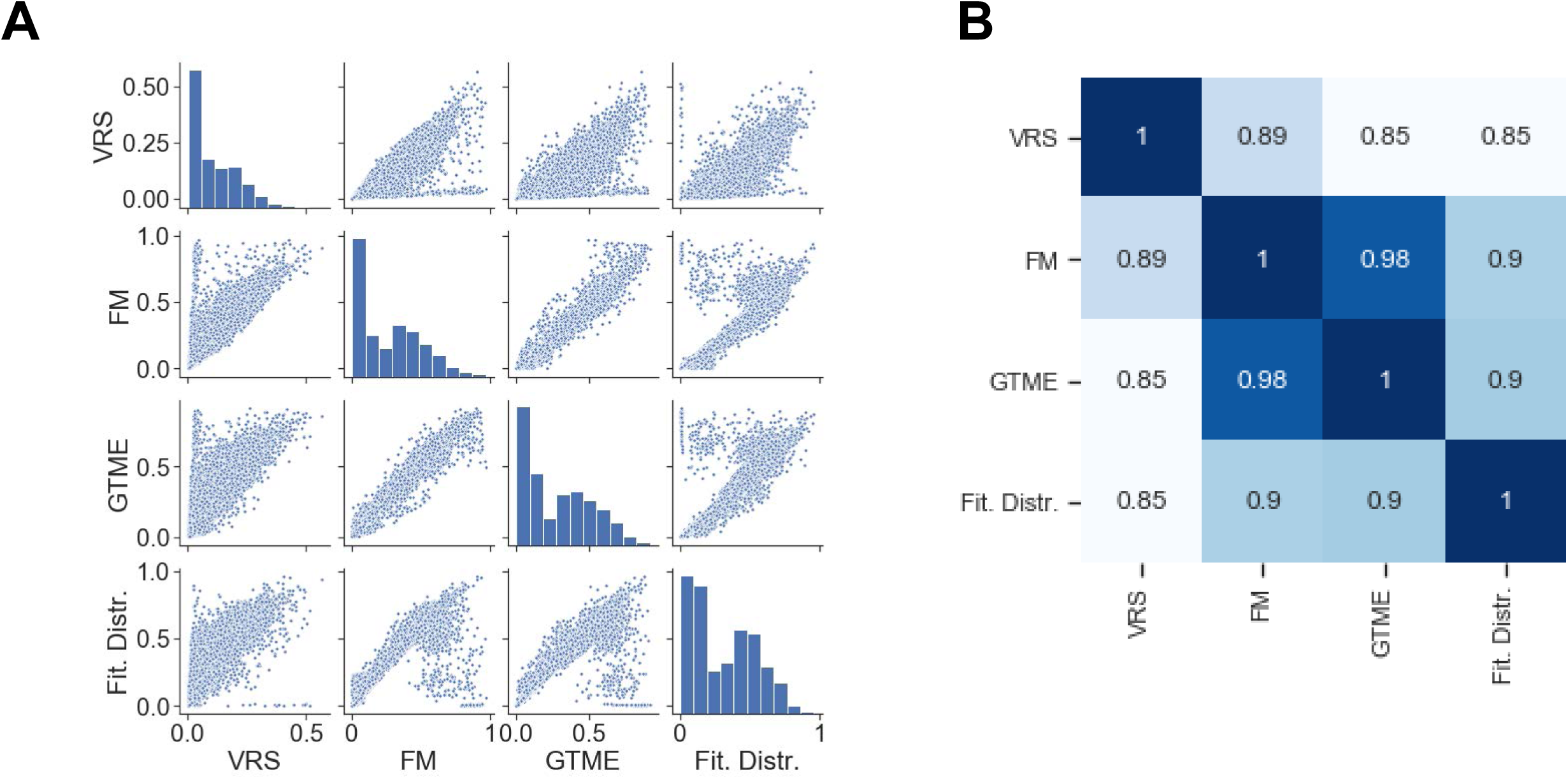
Comparison of dichotomization methods. (A) The pair-wise scatter plot showing the ON cell frequencies of each gene in the dopaminergic neuron dataset, after dichotomization with different methods. (B) The Spearman rank correlation of the ON cell frequencies shown in (A).

**Figure S3.**
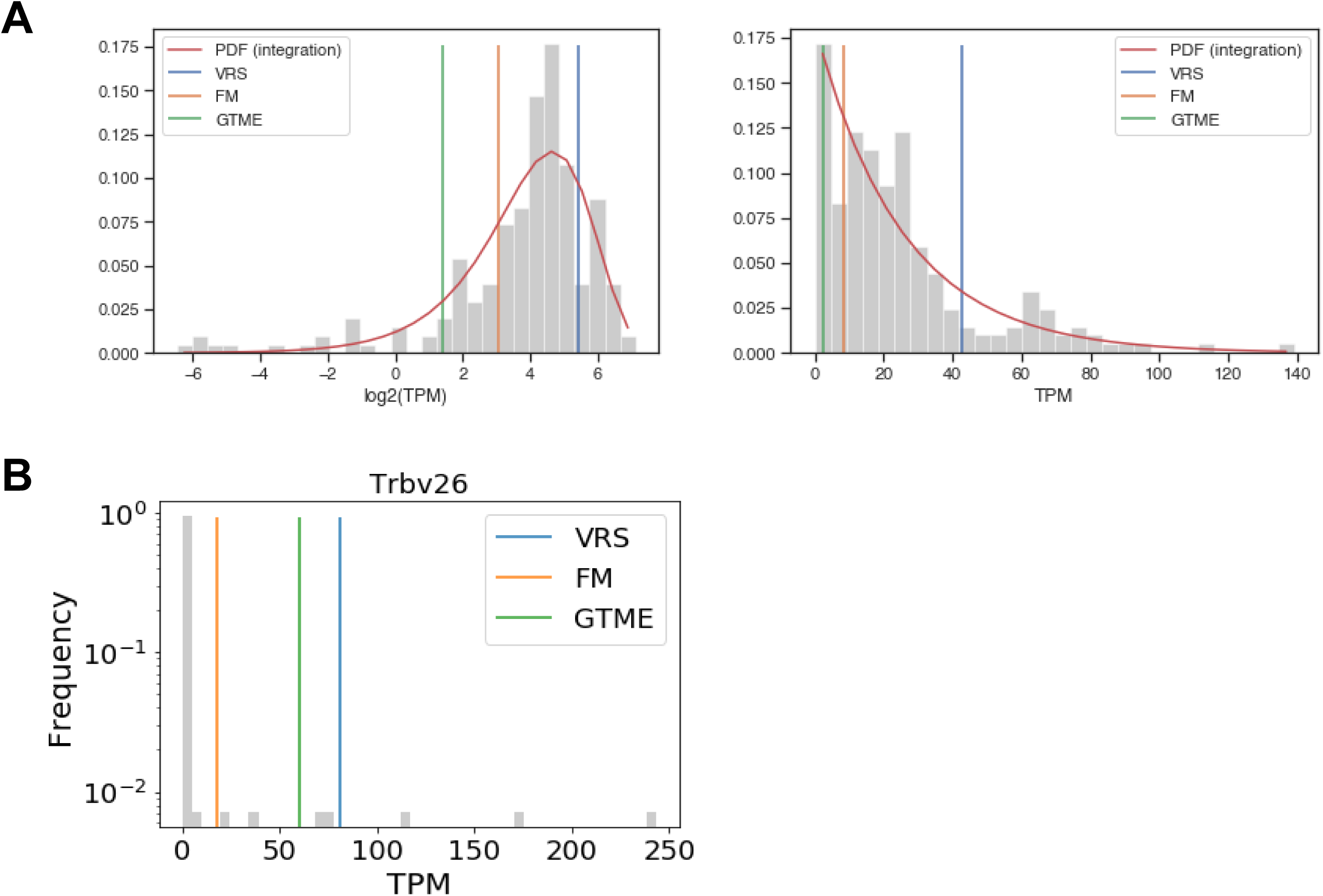
RNA counts fitted with distributions without antimodes. (**A**) The Piezo2 gene is expressed in all cells of the cell population. An exponential function is fitted with lambda = 0.039. The thresholds yield the following ON cell frequencies: 0.17 (VRS), 0.76 (FM) and 0.90 (GTME) TPM. The plot on the right side is the version of the main plat with a linearly scaled x-axis. (**B**) The expression of Trbv26 in Th17 cells. BI = 0.88. The majority of cells do not express this gene and a Poisson distribution with a mean value close to zero is fitted (not shown).

**Figure S4.**
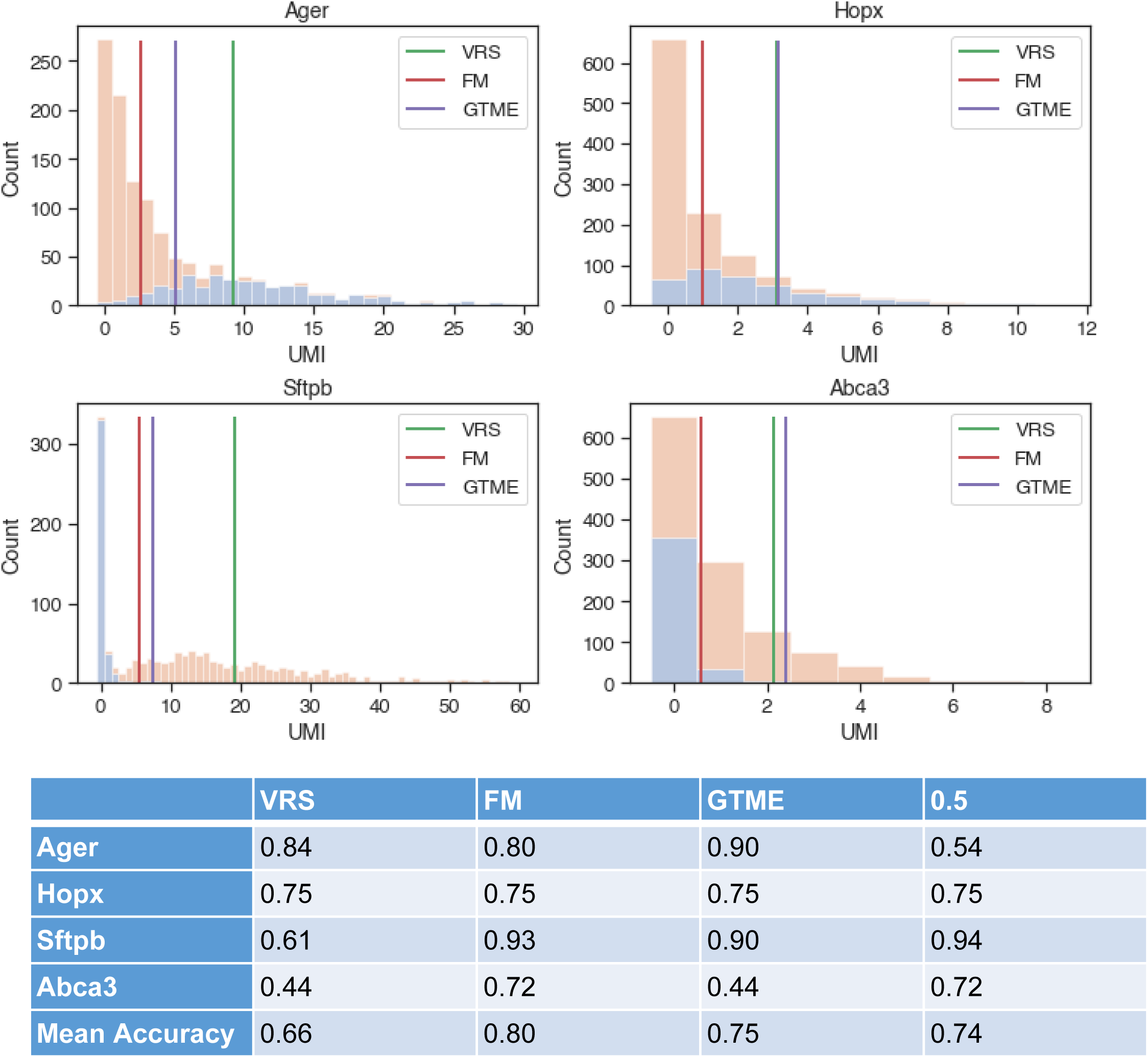
The histograms show expression of four marker genes in Type I (blue) and Type II (orange) alveolar cells. 0.5 denotes a constant threshold at 0.5 UMI, which separates zero from non-zero counts. FM is associated with the lowest misclassification of the two cell types, as evidenced from the highest mean accuracy yielded by this method.

**Figure S5.**
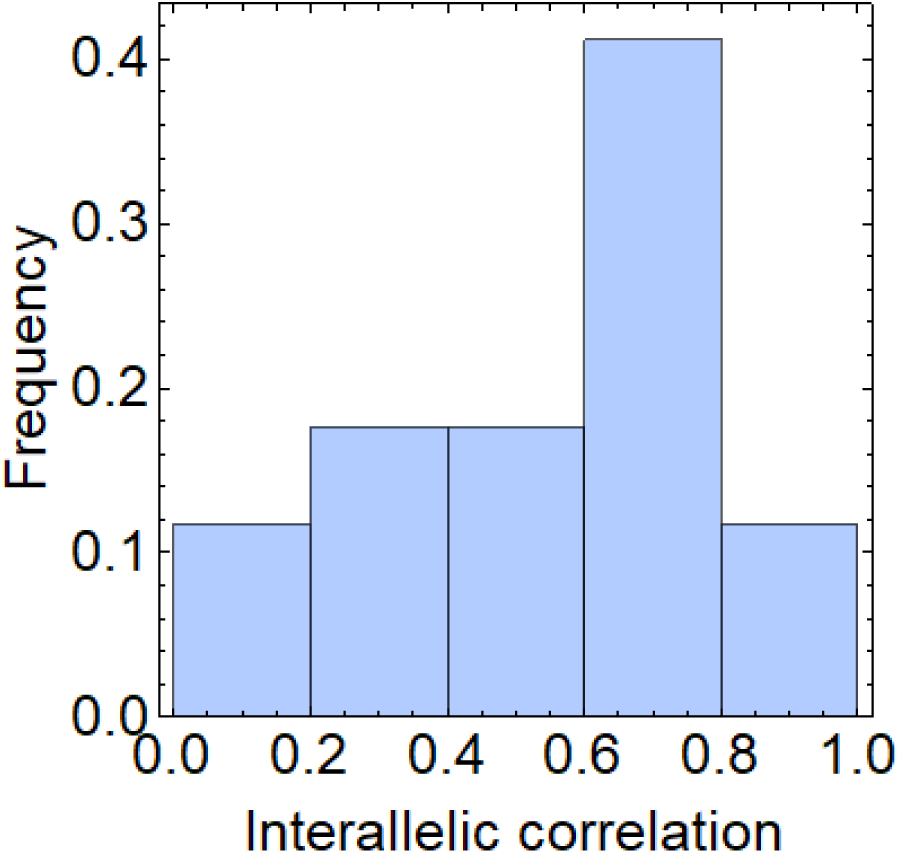
Interallelic correlation in the Pcdh genes expressed in fibroblasts. The beta and gamma isoforms are expressed.

**Table S1.**
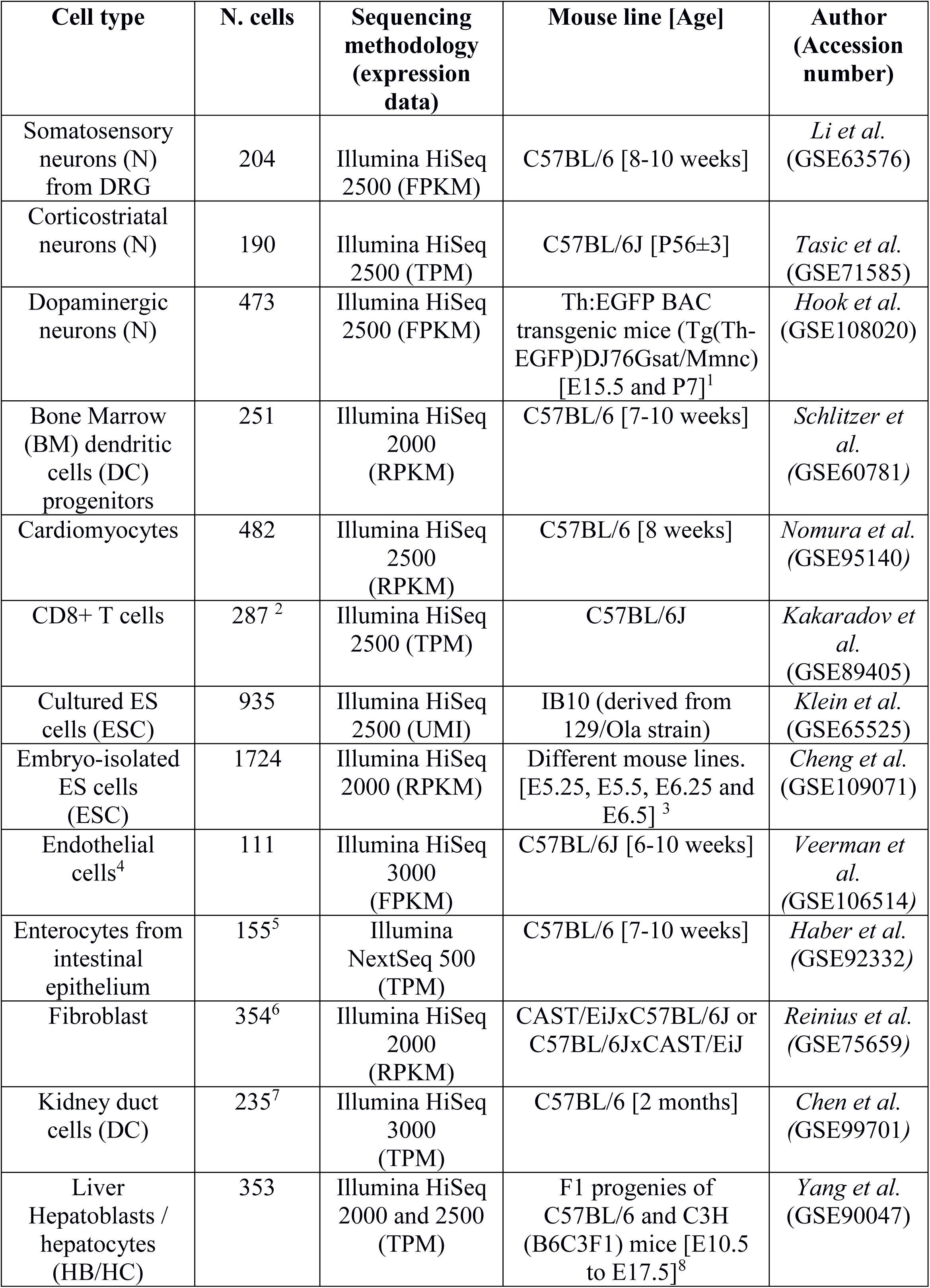
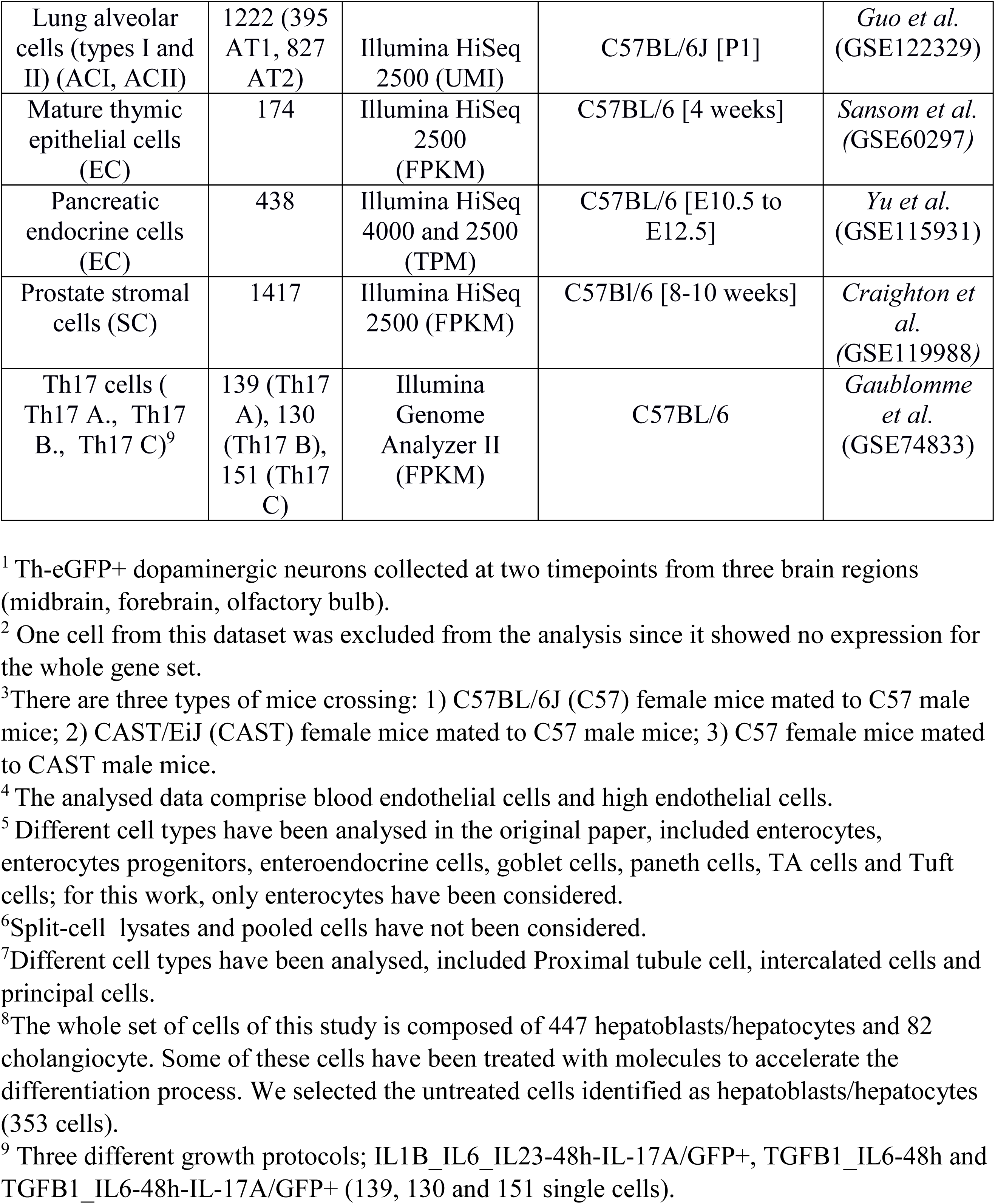
Description of RNA-seq data

**Data S1**. Gene families with stochastic exclusive gene choice. IC values (with 95 confidence intervals) of gene families that belong to the bottom 2.5 percentile of the IC distribution of at least two cell types.

